# Leveraging Crop Wild Relatives and Ploidy Level for Climate-Resilient Annual Ryegrass

**DOI:** 10.1101/2025.10.15.682691

**Authors:** Maria E. Mailhos, Jennifer Timmers, John Erickson, Nicolas Caram, Carlos D. Messina, Esteban F. Rios

## Abstract

Leveraging crop wild relatives and chromosome manipulations are powerful tools for developing climate-resilient agroecosystems. Yet, the combined effects of polyploidy and diverse genetic makeup on plant responses to climate-induced stresses remain generally underexplored, particularly in annual ryegrass (*Lolium multiflorum* Lam.), a major cool-season forage species. In a two-year controlled environment study, we assessed phenotypic responses of a wildtype and the cultivar ‘Marshall’ at diploid (2x = 14) and tetraploid (4x = 28) levels. Plants were grown under 540 and 800 ppm [CO_2_] and at full and 50% evapotranspiration regimes. Anatomical and physiological differences between populations and ploidy levels were limited. Despite genetic background, total biomass production increased by 44% from 540 to 800 ppm [CO_2_], driven by enhanced aboveground growth. While the 2x-wildtype showed a lower leaf-to-stem ratio (a proxy for forage quality) than 2x-Marshall, this gap diminished at the tetraploid condition. These differences, and the lack thereof, highlight the importance of considering both chromosome manipulation and the genetic sourcing of crop wild relatives to expand diversity in cultivated species. Our findings reveal that wild populations can achieve comparable productivity and efficiency to improved populations under climate-change induced environments, while contributing adaptive traits valuable for resilient cropping systems.

**HIGHLIGHT:** Wildtype ryegrass matched improved high-yield cultivar under stress and elevated [CO_2_], suggesting the need for both local adaptation potential and for new breeding targets under climate change.

## INTRODUCTION

The negative impacts of climate change on agroecosystems can be partially offset by a ‘fertilization’ effect in C_3_ species associated with an elevated CO_2_ concentration (e[CO_2_]; Long, 1991; Ainsworth and Rogers, 2007). Increasing temperature reduces CO_2_ diffusivity relative to O_2_ decreasing RuBisCO efficiency and, consequently, carboxylation (Reich et al., 2018; Sollenberger and Kohmann, 2024). However, higher [CO_2_] reduces photorespiration and thus increases net photosynthesis (Drake et al., 1997; Dusenge et al., 2019), counteracting the effects of warming under climate change. Elevated [CO_2_] also improves water-use efficiency via regulation of stomatal conductance and respiration (Drake et al., 1997; Long et al., 2004; Long et al., 2006; Leakey et al., 2009; Leakey et al., 2012; Christy et al., 2018), enhancing photosynthesis rates and, consequently, crop yields (Long, 1991; Morison and Lawlor, 1999; Ainsworth and Rogers, 2007; Leakey et al., 2009; Wang et al., 2012). However, because of diversity in leaf anatomy, stomata size and density, leaf conductance responses to vapor pressure deficits, and canopy architecture, generalizations across species and populations within species are challenging. The genetic regulation of these physiological traits highlights the importance to advance our understanding of how genes, species and populations respond to e[CO_2_], and how these responses affect drought tolerance for developing climate resilient agricultural systems (Toreti et al., 2020).

Crop wild relatives and chromosome manipulations to induce polyploidy are tools and genetic manipulations that could be harnessed in plant breeding to develop cultivars adapted to climate change (Jump et al., 2009). When ploidy level affects stomata size it brings plant functional diversity to the breeding program for the breeder to leverage the photosynthesis response to e[CO_2_] and its consequences on an improved water balance for the crop (Lawson et al., 2010; Doheny-Adams et al., 2012). In general, stomata size is strongly and negatively correlated with stomatal density (Kramer and Boyer, 1995; Wall et al., 2023). Low densities of large stomata lead to lower transpiration rates and, subsequently, greater biomass production in water-limited conditions under e[CO_2_] (Murray, 1995; Doheny-Adams et al., 2012).

Breeders seek to exploit genetic diversity by first assessing local germplasms adapted to specific environmental and stress conditions (Jump and Peñuelas, 2005; Galliart et al., 2019; Kilian et al., 2020; Wall et al., 2023). However, working with genetic diversity is not a simple task, and the difficulty increases with successive cycles of selection (Cooper and Hammer, 1996). The average effect of allelic substitutions within breeding populations are dependent on interactions with other alleles and genomic regions. These regions are selected over time and create a genetic background that shapes context-dependent genetic effects. The opportunity to harness wild relatives for crop improvement depends on the genetic distance between improved cultivars and wild relatives that carry the physiological diversity of interest. Cytogenetic methods, such as polyploidy induction, can rapidly change the genetic context. Hence, combining conventional breeding with cytogenetic tools offers an attractive technique to hasten crop improvement.

Crop varieties that withstand climate-related stresses and are suitable for cultivation in innovative cropping systems will be crucial to maximize risk management, productivity, and profitability under future climate scenarios (Kilian et al., 2020; Wall et al., 2023). However, plant domestication and crop improvement have reduced genetic diversity in most cultivated crops by creating a useful genetic context. It has been argued that this reduction in genetic diversity could limit crop improvement in general and potential adaptation to future challenges (Tanksley and McCouch, 1997; Keneni et al., 2012; Byrne et al., 2018; Swarup et al., 2021). While theoretical sound, empirical evidence supporting or refusing this postulate remains limited. As climate change imposes increasingly complex abiotic and biotic stresses, such as higher temperatures accelerating pests’ evolution, it is imperative to develop genetically resilient varieties to a new range of abiotic and biotic challenges.

Crop improvement depends on the availability and access to novel allelic variants for complex adaptive traits (Jump and Peñuelas, 2005). Crop wild relatives and landraces are valuable sources of these alleles (Cossani and Reynolds, 2015; Seiler et al., 2017), representing heterogeneous, locally adapted versions of domesticated genes that can address current and emerging challenges in stressful environments (Pimentel et al., 1997; Dempewolf et al., 2014; Kilian et al., 2021; Bohra et al., 2022). However, integrating these genetic pools into highly selected cultivars is not trivial. Landraces often exhibit lower or more variable yields and phenology, yet they often possess genes that enhance nutrient uptake and utilization, as well as tolerance to stresses such as drought, salinity, and high temperatures (Keneni et al., 2018). Vast collections of largely unexploited crop genetic diversity are accessible through national and international germplasm banks (Ramirez-Villegas et al., 2022), but these resources remain underprioritized by breeding programs. Barriers include genetic incompatibilities (e.g., the divergence of modern maize breeding pools from landraces, which requires significant breeding efforts to establish compatible genetic backgrounds) and potential trade-offs associated with stress tolerance to a vast array of biotic and abiotic stresses and resource-use efficiency.

Annual ryegrass (*Lolium multiflorum* Lam.) is a C_3_ cool-season species extensively grown in temperate and subtropical regions, including North and South America, Oceania, Asia, and Europe (Rios et al., 2015). It is commonly used for grazing and hay, turfgrass, and more recently as a cover crop (Flower et al., 2012; Yagioka et al., 2015). The species is naturally diploid (2n = 2x = 14), but tetraploid germplasm (2n = 4x = 28) has been developed to improve adaption to specific environments and disease pressures (White and Lemus, 2014). Ryegrasses differing in ploidy level show morphological and anatomical differences (Speckmann et al., 1965; Rios et al., 2015). Tetraploid annual ryegrass tends to have larger leaves, larger stomata, thicker stems, and higher yield than diploid accessions (Rios et al., 2015, 2019). This genetic and ecophysiological diversity makes ryegrasses an excellent case study to investigate how variation in ploidy level can be strategically developed and adopted under different resource availability and environmental conditions.

Prior studies have examined diploid and tetraploid populations of diverse origins around the world (Rios et al., 2015, 2019). Inducing chromosome doubling in diploid ryegrass can generate populations that differ in ploidy level while maintaining the same genetic background. This methodological approach will enable assessing the opportunity to harness ploidy level, at least in ryegrass, to improve crop adaptation to climate change without the confounding of heterogeneous genetic background. The existing genetic diversity of the species provides access to alleles associated with phenology and resilience, traits essential for adaptation to resource-limited environments (Rios et al., 2019; Quesenberry et al., 2022). Yet, how the factorial combination of polyploid ryegrasses with diverse genetic makeup affects anatomical, physiological and functional responses, and how these translate into morphological and productive outcomes under climate-related stresses remains underexplored.

With agricultural intensification, water use efficiency by agricultural cropping systems is an increasing concern further exacerbated by climate variability. The predicted increase in [CO_2_] by the end of the century ranging between 550 and 960 ppm (Masson-Delmotte et al., 2018) is accompanied with spatially and temporally variable rainfall patterns. In some regions, such as southeast United States, the average annual rainfall increased over the past century (Kunkel et al., 2013), but mostly associated to more frequent extreme precipitation events, unlikely overcoming drought events during the year. In this scenario, it is imperative to understand how genetic diversity including ploidy level of annual ryegrasses ameliorate drought events and respond to e[CO_2_] to develop climate resilient agroecosystems. Notably, most studies assessing the impacts of e[CO_2_] on plant responses have considered levels between 550 and 700 ppm. Yet, crop responses are less clear growing under concentrations above 800 ppm, a plausible scenario by 2100. The objectives of the study were to examine the combined effect of cytogenetics, through plant ploidy level (2x vs. 4x), and germplasm diversity (bred vs. wildtype) on anatomical, physiological and morphological responses of annual ryegrass under two e[CO_2_] (540 and 800 ppm) and two water availability levels (50% and 100% of potential evapotranspiration).

## MATERIALS AND METHODS

### Experimental Design

A controlled-environment experiment was conducted at the University of Florida Climate Change Greenhouse Facility in Gainesville, Florida (29.39 N, 83.40 W). The greenhouses were controlled for temperature, atmospheric [CO_2_] and relative humidity conditions (Allen et al., 2018). The experiment consisted in two runs performed in two years. The first run of the experiment was conducted from November 2016 to February 2017 and the second run was conducted from November 2017 to February 2018.

The experiment was arranged as a randomized complete block design with four replicates (r = 4) and a split-plot restriction on randomization. The first factor was the atmospheric [CO_2_] which consisted in two levels of 540 (±1.1) ppm and 800 (±0.5) ppm, chosen based on the lower and upper ends of the IPCC [CO_2_] stabilization scenarios (IPCC 2018). The experimental design had a split-plot restriction on randomization where the atmospheric [CO_2_] was randomly assigned at the main plot level (n=4). Thus, eight greenhouses were the main plots, where four were managed under 540 ppm and four under 800 ppm. At the subplot level, irrigation (full evapotranspiration (ET) versus 50% ET), ploidy level (diploid vs. tetraploid), and population (wildtype vs. bred type) were randomly assigned to the experimental units, corresponding to 3.2-L pots. The irrigation levels were thought to simulate scenarios of water-limited and ‘unlimited’ conditions. The two populations (wildtype vs. bred type) were selected from the University of Florida Ryegrass Breeding program, the cultivar Marshall and the plant introduction 241586 obtained from the U.S. National Plant Germplasm System.

For tetraploid induction, germinating seeds from both diploid populations were treated with 0.5% colchicine [(S)-N-(5,6,7,9-tetrahydro-1,2,3,10-tetramethoxy-9-oxobenzo [a]heptalen-7-yl) acetamide; Sigma Aldrich, Darmstadt, Germany] for three hours (Figure 1). Seedlings were rinsed with purified water, placed on Petri dishes with filter paper for two days, then individual plants were transplanted into 7.5 cm pots. Plants were conditioned in high humidity for several days before being transferred to the greenhouse. Tetraploid selection was performed using flow cytometry following Rios et al. (2015). Five weeks after colchicine treatment, leaf samples were collected from at least three tillers per pot, cut into 0.5 to 1.0-mm segments, and placed in a Petri dish with 500 μl extraction buffer (CyStain PI absolute P, Partec GmbH, Münster). The leaf tissue was chopped for 30 seconds with a razor blade, incubated on ice for 30 seconds in the extraction buffer, filtered through Partec 50 μm CellTrics (Partec GmbH, Münster), and stained with 2 ml of propidium iodide and RNase (CyStain PI absolute P, Partec GmbH, Münster). Samples were incubated on ice for at least 30 minutes and analyzed with the BD Accuri C6 Flow Cytometer using the FL2-A channel (Accuri Cytometers, Ann Arbor, MI) at the University of Florida Interdisciplinary Center for Biotechnology Research, Gainesville, FL. The tetraploid populations were advanced two generations to produce enough seed for the trials. Each generation involved polycrosses between 20 to 30 plants, with populations isolated to prevent pollen flow between the wild-type and Marshall. Both diploid lines and the resultant tetraploid lines were included in this study. Each combination of population x ploidy level x irrigation was replicated in three pots per greenhouse.

**Figure 1.**
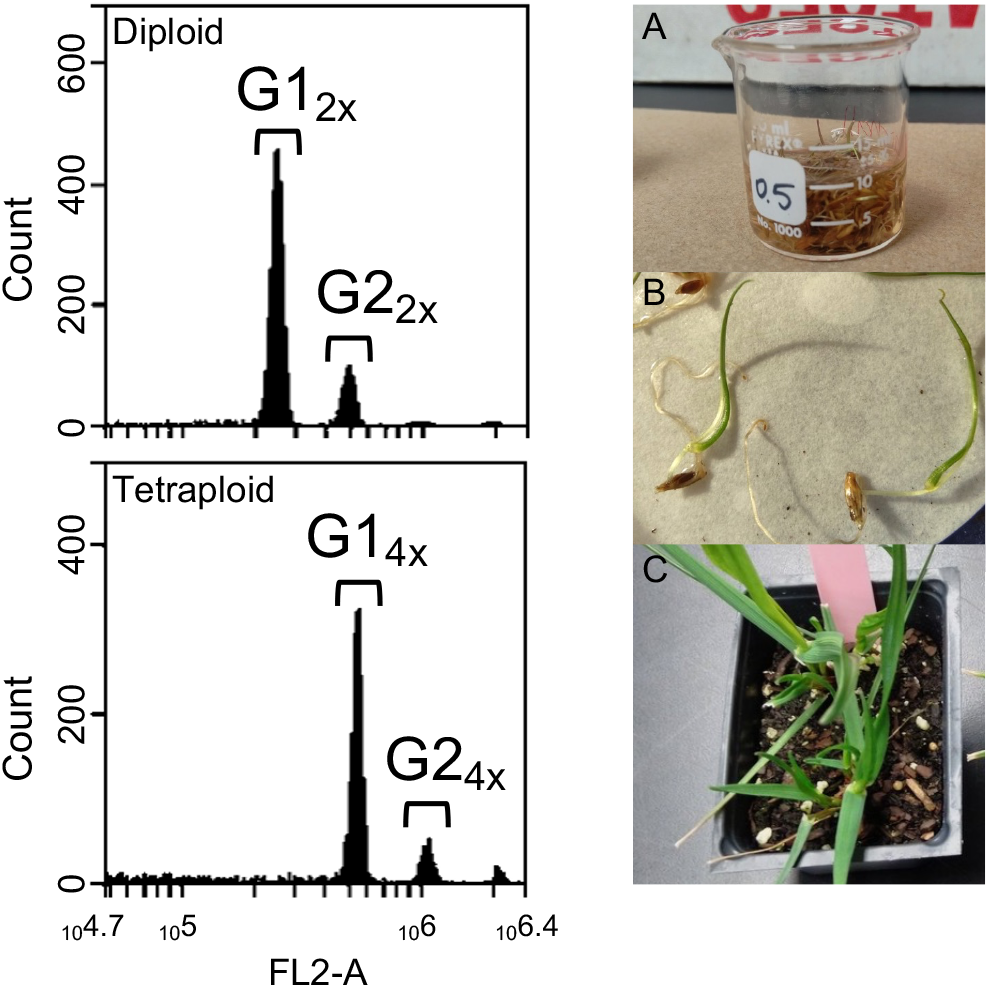
Flow cytometry analysis (left panels) and colchicine treatment procedure (right panels) for induction of tetraploidy in annual ryegrass (*Lolium multiflorum*). Left panels: Histograms show fluorescence intensity of propidium iodide bound to DNA in cell nuclei extracted from leaves of diploid (top) and colchicine-induced tetraploid (bottom) genotypes. Peaks G1_2_x and G2_2_x correspond to diploid nuclei in G1 and G2 phases of the cell cycle, while G1_4_x and G2_4_x represent tetraploid nuclei. The x-axis indicates fluorescence intensity (logarithmic scale), proportional to DNA content; the y-axis shows the number of nuclei analyzed. An internal standard was not used; samples were compared against untreated diploid controls (2n = 2x = 14) for the Marshall cultivar and wild type. Right panels (A–C): A) Approximately 40 germinating seeds were submerged in 0.5% colchicine solution for 3 h. B) Treated seedlings were rinsed with purified water and incubated on filter paper in Petri dishes for 2 days. C) Seedlings showing abnormal growth (leaf curliness, shorter and thicker leaves, darker green color) were transplanted into 7.5 cm pots, maintained under high humidity for several days, and later transferred to the greenhouse for ploidy assessment using flow cytometry.

Seeds from each population and ploidy were germinated for the study. Approximately two weeks after germination, seedlings of uniform size were transferred to 3.2-L pots containing a root zone mix made from 10% (by volume) potting mix (Sungro Professional Growing Mix) and 90% coarse-textured sand. Thirty-two pots were randomly placed on tables in each climate-controlled polycarbonate greenhouse room ([CO_2_] whole plot). The air temperature in each room was controlled with a square wave day/night temperature function of 25/15 °C. From 09:00 to 21:00 temperature was set to 25 °C and from 21:00 to 09:00 it was set to 15 °C. No supplemental lighting was used. Relative humidity was maintained at 60% in each room during the experimental period. Plants received a single application of fertilizer at a rate of 50 kg nitrogen (N) ha^-1^ using Peter’s Excel® 15-5-15 Cal-Mag special (11.8% nitrate N [NO_3_-N] 1.1% ammoniacal N [NH_4_-N], and 2.1% urea N) after approximately 45 days of growth. Plants were irrigated by a drip irrigation system set up on timer. The drip irrigation system was calibrated to supply either at a full ET rate, based on average daily weight loss in the 540 ppm (n=10) and 800 ppm rooms (n=10) or 50% ET rate (half the water supplied of the full ET treatment).

### Data Collection

After approximately 90 days of vegetative growth and just prior to harvest, leaf gas exchange data were collected using a Li-Cor (Li-Cor Inc., Lincoln, NE) 6400 XT Portable Photosynthesis System. Stomatal conductance (g_s_; mol H_2_O m^-2^ s^-1^), *A/Ci* curves (photosynthesis rate, *A*, versus internal CO_2_ concentration, *Ci*) and light-saturated photosynthesis (*Asat*; μmol CO_2_ m^-2^ s^-1^) were measured on the newest fully expanded leaves between 10:00 h and 15:00 h during sunny sky conditions. The maximum rate of RuBisCO carboxylation (V_cmax_) was calculated from Farquhar et al. (1980) model as described in Sharkey (2015). *A/Ci* curves were taken on three sampling units (pots) within greenhouse rooms (n=24). The leaf chamber environment was set to a photosynthetic photon flux density of 1200 µmol m^-2^ s^-1^, using the 6400-02 light emitting diode light source, a flow rate of 500 µmol s^-1^, a relative humidity of approximately 60%, and a block temperature of 25 °C. The *A/Ci* curves included nine chamber [CO_2_] set points ranging from 50 to 1000 ppm for the 540-ppm treatment, and to 1200 ppm for the 800-ppm treatment. The water use efficiency (WUE; μmol CO_2_ mol H_2_O ^-1^) was estimated as the ratio between photosynthesis rate, *A*, and stomatal conductance, g_s_).

Once *A/Ci* data was collected, an epidermal impression was taken using nail polish of the abaxial leaf surface of a fully emerged, healthy leaf. The impression was placed on a slide and guard cell measurements were taken using an ocular micrometer at 40x magnification. Photographs were taken using the v. 6.2.0 INFINITY Camera and Software® (Lumenera Corporation, 7 Capella Court, Ottawa, ON, Canada).

Plants were then harvested for leaf blade, stem, and root dry matter (DM) content. The whole plant was cut at soil level. Tillers and number of leaves per tiller were counted and recorded. Leaf blades were removed from the stems at the collar region. Leaf blades and stems were dried at 60 °C for 72 hours. The potting medium was shaken loose from roots. Remaining debris was washed off from the roots and then were dried at 60 °C for 72 hours. Leaf blades, stems, and roots were weighed, and biomass measurements recorded.

### Data Analysis

Linear mixed models were performed to analyze anatomical, physiological and morphological responses using the *lmer* function of the ‘lme4’ package (Bates et al., 2015) in R 4.4.0 (R Development Core Team). The [CO_2_] (540 vs. 800 ppm), irrigation (50 vs. 100% ET), population (Marshall vs. PI 241568) and ploidy (2x vs. 4x) were included as fixed effects, while block was included as random effect. Runs (years) were included as single effect factor in the model to capture potential differences between runs that could mask differences between other factors. Differences between means were studied using emmeans function of the emmeans package (Lenth et al., 2018). Statistical differences were declared at *P* < 0.05. Residuals were studied using DHARMa package (Hartig and Hartig, 2017). To explore potential associations between stomatal length and physiological responses (photosynthesis rate, stomatal conductance and maximum rate of RuBisCO carboxylation) a Pearson correlation test was performed between these responses (*P* < 0.05).

## RESULTS

### Anatomical and physiological responses

Plants showed different anatomical and physiological responses to the various factors examined at the end of the growing season. Stomata were 30% longer in tetraploid and 13% longer in wildtype ryegrasses compared to diploid and cultivated types, respectively (Figure 2A), while other factors showed no significant effect. Under 540 ppm, ploidy was the main driver of photosynthesis rate differences, where diploids showed 64% higher rates per leaf area than tetraploids. In contrast under 800 ppm, irrigation became the dominant factor, where photosynthesis rate per area was surprisingly 102% higher under 50% than 100% ET (Figure 2B) at this single pre-harvest measurement. A similar pattern was observed for the maximum RuBisCO carboxylation rate (V_cmax_) in tetraploids, which was 54% higher under 50% compared to 100% ET (39.4 vs 25.6 ±2.5 μmol m^-2^ s^-1^). Such differences were not observed for diploid populations, averaging 30.6 μmol m^-2^ s^-1^ across irrigation levels (Figure 2C).

**Figure 2.**
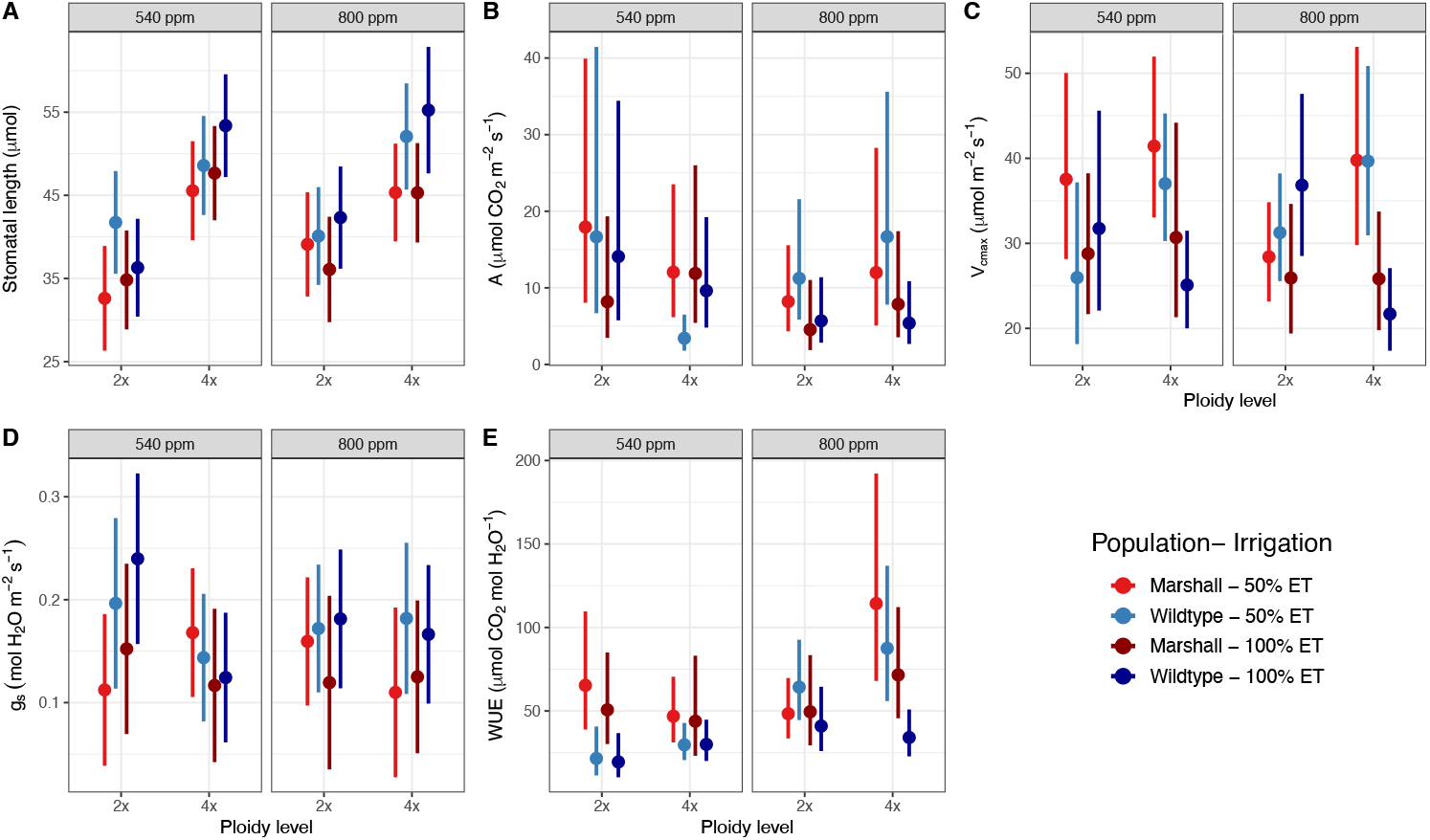
Anatomical and physiological responses of the two populations of ryegrass varying in ploidy level to different environments. Populations are diploid (2x) and tetraploid (4x) Marshall (bred type) and wildtype (PI 241568) ryegrasses, growing under 540 and 800 ppm of [CO_2_] and with full irrigation (100% ET) and 50% of ET. Vertical error bars denote the 95% confidence interval. (**A**) is the stomatal length; (**B**) is photosynthesis rate (*A*); (**C**) is the maximum rate of RuBisCO carboxylation (V_cmax_); (**D**) is the stomatal conductance (g_s_); and (**E**) is the water use efficiency (WUE).

At this single point, stomatal conductance was 29% higher in the wildtype than in Marshall (0.168 vs 0.130 ±0.019 mol H_2_O m^-2^ s^-1^; Figure 2D). This difference was largely driven by a 63% increased stomatal conductance in the diploid wildtype under 540 ppm compared to other ploidy-population combinations. This difference was no longer detected in plants growing under 800 ppm. The higher stomatal conductance of the wildtype at 540 ppm corresponded to lower water use efficiency, i.e., the ratio between photosynthesis to stomatal conductance, whereas at 800 ppm, tetraploid Marshall showed 85% higher water-use efficiency than diploids (Figure 2E). No significant associations were found between stomatal length and physiological traits at the end of the growing season (*r* = -0.13 to 0.16; *P* > 0.30).

### Morphological responses

The limited differences in anatomical, and especially physiological responses between populations and ploidy levels at the end of the growing season were reflected in cumulative biomass production. Across populations, ploidy, and water levels (*P* > 0.25), total biomass was 44% greater under 800 ppm than 540 ppm (4.03 vs. 2.79 ±0.33 g pot^-1^). This indicates that the wildtype performed as well as the cultivated type under varying growth condition and stress levels. This increase in total production under 800 ppm was driven by aboveground production (2.29 vs. 1.60 ±0.17 g pot^-1^; +43%, Figure 3), while root production remained similar across treatments (1.46 ±0.14 g pot^-1^). Consequently, the aboveground-to-belowground biomass ratio was higher under 800 ppm than 540 ppm (1.43 vs. 1.17 ±0.09). No evidence suggested that ploidy or population affected total or partitioned biomass.

**Figure 3.**
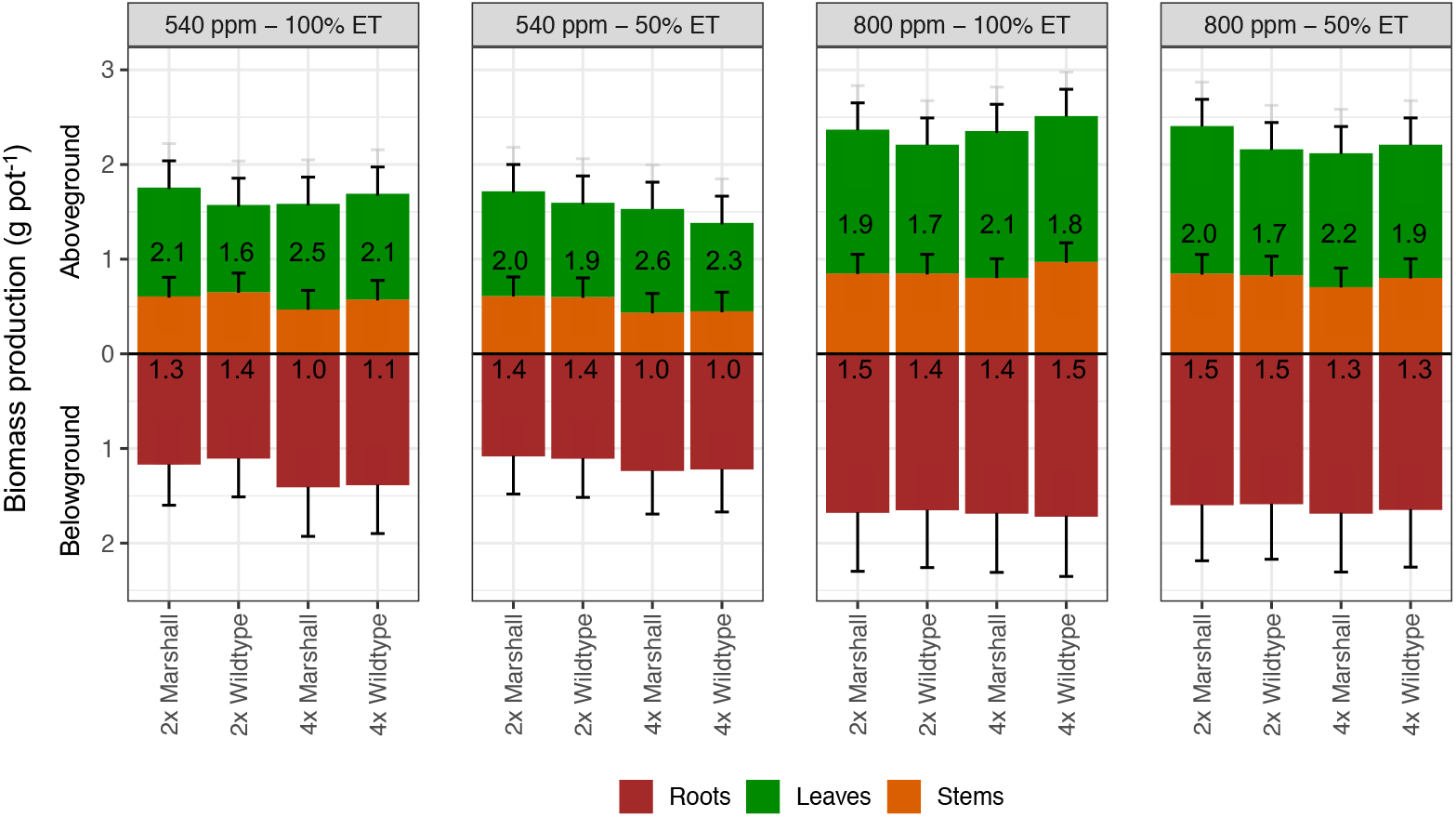
Biomass production of the two populations of ryegrass varying in ploidy level to different environments. Populations are diploid (2x) and tetraploid (4x) Marshall (bred type) and wildtype (PI 241568) ryegrasses, growing under 540 and 800 ppm of [CO_2_] and with full irrigation (100% ET) and 50% of ET. Belowground biomass production denotes the root production (brown), and aboveground biomass production denotes the stems (orange) and leaves (green) biomass production. Grey vertical error bars at the top of the green bar denote the total aboveground biomass production (stems + leaves) 95% confidence intervals. Black vertical error bars denote the 95% confidence intervals for the individual components (roots, stems, leaves). Numbers on the green bars depict the leaf-to-stem ratio for each combination of ploidy level and population under 540 and 800 ppm, *SE ± 0*.*16*. Numbers on the brown bars depict the aboveground-to-belowground ratio for each combination of ploidy level and population under 540 and 800 ppm, *SE ± 0*.*1*.

Aboveground partitioning between stems and leaves, a trait of great agronomic importance due to its link with forage nutritive value, revealed that plants produced more leaves and stems under 800 ppm than 540 ppm (1.46 vs. 1.05 ±0.10 g pot^-1^ and 0.83 vs. 0.55 ±0.07 g pot^-1^, respectively, Figure 3). Marshall produced 9% more leaf biomass than the wildtype (1.31 vs. 1.20 ±0.08 g pot^-1^), resulting in a larger leaf-to-stem ratio (2.18 vs. 1.87 ±0.08). Tetraploids produced less stem biomass than diploids under 540 ppm (0.48 vs. 0.62 ±0.08 g pot^-1^), associated with a lower tiller number (6.6 vs. 8.8 tillers pot^-1^). e[CO_2_] increased tiller number (9.2 vs. 6.3 ±0.5 tiller pot^-1^ under 800 vs. 540 ppm), explaining the lower leaf-to-stem ratio of tetraploids at 800 ppm compared to 540 ppm (1.99 vs. 2.38 ±0.12); diploids averaged 1.87 ±0.12. Notably, leaf number per tiller was consistent across populations, ploidy levels and environmental conditions, averaging 4.4 ±0.13 leaves per tiller.

## DISCUSSION

Developing new resilient genotypes to face uncertain climate change scenarios is relevant for future food systems (Kilian et al., 2021; Bohra et al., 2022). In this regard, exploiting crop wild relatives and polyploidy provides alternatives for adapting farming systems to stressful and resource-limited environments (Pimentel et al., 1997; Dempewolf et al., 2014). Despite the relevance of annual ryegrass in global food systems, the potential use of wild relatives and variation in ploidy level for adaptation to limiting environments remains underexplored (Rios et al., 2019; Quesenberry et al., 2022). Furthermore, the broad ecophysiological and genetic diversity of ryegrass provides an ideal framework to explore how ploidy variation can be leveraged to optimize performance across resource and environmental gradients. In this controlled-environment study, we found a promising use of a wildtype ryegrass in producing biomass as efficiently as a high-yield released genotype under water-limited conditions and e[CO_2_]. This reveals the potential of wildtype populations on achieving production efficiency comparable to improved genotypes, while potentially carrying alleles that enhance physiological resilience and local adaptation. Given these findings, further research could test the hypothesis that genetic distance between improved cultivars and wild relatives determines opportunities to harness adaptive alleles from landraces and relatives, as previously proposed (Reynolds et al., 2007).

Differences in anatomical and physiological responses at a single time point between populations and ploidy levels did not translate into major morphological responses. Nonetheless, some contrasting patterns provide insight into underlying mechanisms. For instance, both populations produced similar total, aboveground and belowground biomass under comparable growth conditions, but plants grown at 800 ppm CO_2_ achieved higher biomass production, reflecting the well-documented CO_2_ fertilization effect (Drake et al., 1997; Dusenge et al., 2019), and coinciding with generally higher photosynthetic rates and WUE per area (Figure 2). e[CO_2_] typically increases carboxylation efficiency and induces partial stomatal closure, increasing transpiration efficiency and often resulting in increased growth and DM yields (Assman, 1993; Vavasseur and Raghavendra, 2005; Lawson, 2009). However, under certain conditions, Marshall showed higher instantaneous WUE than the wildtype, indicating a potential target for improving the wildtype adaptation to resource-limited environments.

The difference in WUE between populations could reflect multiple factors. Marshall was initially selected and bred for increased winter hardiness (Arnold et al., 1981), which may have co-selected linked genes for traits promoting increased WUE. Additionally, selection for cold tolerance may have also favored alleles involved in drought tolerance, such as DREB (Dehydration-Responsive Element Binding) transcription factors. These regulate the expression of stress-responsive genes, enhancing tolerance to drought, cold and other abiotic stresses (Agarwal et al., 2006). Consequently, selection for cold tolerance in Marshall may have indirectly selected for drought resilience and improved WUE. Further, while few studies have assessed genetic correlations between phenotypic traits and WUE, breeding populations of *Arabidopsis thaliana* (L.) Heynh. have shown improvements in WUE, unlike wild-type populations, suggesting that selection for traits such as vegetative growth and reproductive performance (*vegetative growth = WUE × total water transpired*; *yield = vegetative growth × harvest index*) can indirectly enhance WUE (Edwards et al., 2012).

The absence of clear associations between stomatal length and physiological responses at the end of the growing season likely reflects compensatory mechanisms at the leaf level and the inherent limitations of evaluating a single anatomical trait at one time point. Stomatal size and density determine total pore area per unit of leaf area (Franks and Beerling, 2009), and tetraploids typically have larger stomata than diploids (Rios et al., 2015; Figure 2A). Despite the well-established relationships between stomatal anatomy and instantaneous gas exchange per unit of leaf area, the lack of association observed in our study likely results from the timing of measurements, taken once during the growing season, just before harvest, the relatively narrow range of [CO_2_] treatments, and other growing conditions constraints. Further, stomatal behavior rather than size alone largely determines gas exchange, as dynamic regulation of stomatal aperture can buffer intergenotypic differences in physiological responses (Lawson and Blatt, 2014).

The low assimilation rates observed across factors by the end of the growing season (Figure 2B) suggest that plants operated below their potential biochemical capacity at this stage. Yet, the absence of data on stomatal density limits interpretation of potential diffusion constraints, particularly in tetraploids, which showed longer stomata and may therefore have lower stomatal densities, restricting CO_2_ uptake. Although tetraploidy did not enhance instantaneous physiological responses, it may have improved regulatory stability and altered biomass partitioning (Figure 3). Tetraploids produced a greater proportion of leaves relative to stems, indicating a shift toward metabolically richer tissues that could enhance light interception and energy return per unit of carbon invested (Wilson et al., 1999). Such allocation patterns may help sustain productivity and forage quality despite lower area-based assimilation rates and emphasize the importance of considering the energic concentration of assimilated tissues when evaluating plant responses (Caram et al., 2025).

This shift in allocation connects directly to a key agronomic trait: the proportion of leaves in total biomass production. Leaf proportion is particularly relevant in forage species because leaves are more nutritious and preferred in livestock systems (Rios et al., 2015). Therefore, ‘leafy’ ryegrasses are generally more desirable than ‘stemmy’ ones. In our study, Marshall showed a higher leaf-to-stem ratio than the wildtype, although this difference diminished at 800 ppm compared with 540 ppm. Tetraploids consistently showed a higher leaf-to-stem ratio with fewer tillers compared to diploids, as also reported by Rios et al. (2015). These findings suggest that polyploidy may promote shifts in biomass allocation toward leaves, improving forage quality while maintaining yield potential (Rios et al., 2019; Quesenberry et al., 2022). The attenuation of ploidy-related differences at 800 ppm CO_2_ further supports the idea that CO_2_ enrichment reduces trait divergence by alleviating water limitations and enhancing overall C gain.

Finally, the convergence in aboveground and belowground allocation patterns among populations under 800 ppm suggests that genetic and cytogenetic diversity did not compromise root function. The wildtype allocated assimilates to roots comparably to the cultivated genotype, which could support nutrient uptake and sustained productivity under stress. Together, these results indicate that combining wild genetic backgrounds with induced polyploidy could enhance physiological stability and resource-use efficiency in ryegrass. This reinforces the need to explore crop wild relatives and landraces adapted to local and limited environments to meet the challenges of farming in stressful environments (Pimentel et al., 1997; Dempewolf et al., 2014; Kilian et al., 2021; Bohra et al., 2022). Future studies integrating long-term physiological measurements, stomatal density, mesophyll conductance, and C allocation dynamics would help disentangle the relative contribution of anatomical versus biochemical mechanisms in shaping adaptation to e[CO_2_] and drought, thereby, contributing to the design of climate resilient cropping systems.

## Abbreviations

[CO_2_]: CO_2_ concentration;
*A*: photosynthesis rate;
*Ci*: internal CO_2_ concentration
e[CO_2_]: elevated CO_2_ concentration;
ET: evapotranspiration;
g_s_: stomatal conductance;
V_cmax_: maximum RuBisCO carboxylation rate;
WUE: water use efficiency.

## SUPPLEMENTARY DATA

No supplementary data is provided.

## AUTHOR CONTRIBUTIONS

JT, JE and ER: conceptualization, methodology and investigation; NC: formal analysis; MM and NC writing – original draft preparation; JT, JE, MM, NC, CM and ER writing – review & editing; ER: supervision; ER: funding acquisition.

## DATA AVAILABILITY

Data generated during this study are available from the corresponding author on reasonable request.

